# *In silico* analysis of *CDC73* gene revealing 11 novel SNPs associated with Jaw Tumor Syndrome

**DOI:** 10.1101/729764

**Authors:** Abdelrahman H. Abdelmoneim, Alaa I. Mohammed, Esraa O. Gadim, Mayada A.Mohammed, Sara H. Hamza, Sara A. Mirghani, Thwayba A. Mahmoud, Mohamed A. Hassan

## Abstract

**Back ground:** hyperparathyroidism-jaw tumor (HPT-JT) is an autosomal dominant disorder with variable expression, with an estimated prevalence of 6.7 per 1,000 population. Genetic testing for predisposing CDC73 (HRPT2) mutations has been an important clinical advance, aimed at early detection and/or treatment to prevent advanced disease. The aim of this study is to assess the effect of SNPs on *CDC73* structure and function using different bioinformatics tools.

**Method:** Computational analysis using eight different *in-silico* tools including SIFT, PROVEAN, PolyPhen-2, SNAP2, PhD-SNP, SNPs&GO, PMut and Imutant were used to identify the impact on the structure and/or function of *CDC73* gene that might be causing jaw tumour.

**Results:** From (733) SNPs identified in the *CDC73* gene we found that only Eleven were deleterious to the function and structure of protein and expected to cause syndrome.

**Conclusion:** Eleven substantial genetic/molecular aberrations in *CDC73* gene were identified that could serve as actionable targets for chemotherapeutic intervention in patients whose disease is no longer surgically curable.

## 1. Introduction

Primary hyperparathyroidism (PHPT) is a disease caused by the autonomous over-secretion of parathyroid hormone (PTH) due to adenoma and hyperplasia of the parathyroid gland, with an estimated prevalence of 6.7 per “1,000” population. The majority of PHPT cases are not inherited, and some are caused by genetic mutations, including multiple endocrine neoplasia (MEN), familial hypocalciuric hypercalcemia, neonatal severe hyperparathyroidism, familial isolated hyperparathyroidism and hyperparathyroidism-jaw tumor (HPT-JT) syndrome (caused by mutations in the CDC73 gene).[1–5]

HPT-JT (OMIM #145001) was first observed by Jackson in 195. It is an autosomal dominant disorder with variable expression.[6–8] It includes parathyroid adenomas, fibro-osseous jaw tumors, uterine tumors and renal diseases such as hamartomas, polycystic disease and Wilms tumors or adenocarcinoma. Diagnosis of HPT-JT is important because of its genetic involvement and 24% malignant transformation.[9, 10] The nuclear medicine imaging - especially the scintigraphy parathyroid with 99m Tc-MIBI (methoxyisobutyl-isonitrile)-has an important role in outlining the diagnosis.[11] It has been estimated that approximately 70% of patients affected by this mutation may develop PHPT.[12]

CDC73-related (HPT-JT) syndrome results from truncating (80%) or missense variants in the *CDC73* gene (also known as HRPT-2), which encodes the parafibromin protein.[13–16]

This study is unique because it is the first insilico analysis of *CDC73* gene associating it with jaw tumor syndrome. The aim of this study is to assess the effect of SNPs on CDC73 structure and function using different bioinformatics tools.

## 2. Methodology

### Source of retrieving nsSNPs

The SNPs related to the human gene *CDC37* were obtained from single nucleotide database (dbSNP) in the National Center for Biotechnology Information (NCBI) web site. www.ncbi.nlm.nih.gov. And the protein sequence was obtained from UniProt database. www.uniprot.org.

### Severs used for identifying the most damaging and disease related SNPs

#### ❖ Four server used for assessing the functional impact of deleterious nsSNP

##### 1 SIFT (Sorting intolerant from tolerant)

SIFT is the first online server that was used in our assessment. It predicts whether an amino acid substitution affects protein function based on sequence homology and the physical properties of amino acids. Sift score range from 0 to 1. The amino acid substitution is predicted **deleterious** if the score is ≤0.05, and **tolerated** if the score is ≥ 0.05. The protein sequence that obtained from UniProt and the substitutions of interest were submitted to SIFT server, then according to the score the substitution will be either **deleterious** or **tolerated**. The deleterious SNPS were further evaluated [17]. It is available online at http://www.sift.bii.a-star.edu.sg/

##### 2 Polyphen2 (prediction of functional effect of human nsSNPs)

PolyPhen-2 (Polymorphism Phenotyping v2) it is another online server, available as software and via a Web server predicts the possible impact of amino acid substitutions on the stability and function of human proteins using structural and comparative evolutionary considerations. It performs functional annotation of single-nucleotide polymorphisms (SNPs), maps coding SNPs to gene transcripts, extracts protein sequence annotations and structural attributes and builds conservation profiles. It then estimates the probability of the missense mutation being damaging based on a combination of all these properties. PolyPhen-2 features include a high-quality multiple protein sequence alignment pipeline and a prediction method employing machine-learning classification.[18]

The output of the PolyPhen-2 prediction pipeline is a prediction of probably damaging, possibly damaging, or benign, along with a numerical score ranging from 0.0 (benign) to 1.0 (damaging). This three predictions means that, when the prediction is "probably damaging" indicates damaging with high confidence, "possibly damaging" indicates damaging with low confidence, and "benign" means that the query substitution is predicted to be benign with high confidence. It is available online at http://genetics.bwh.harvard.edu/pph2/. Only the damaging SNPs were further evaluated.

##### 3 PROVEN (protein variation effect analyzer)

Was the third software tool used which predicts whether an amino acid substitution has an impact on the biological function of a protein. PROVEAN is useful for filtering sequence variants to identify nonsynonymous or indel variants that are predicted to be functionally important. The result obtained from this web site is either deleterious when the score is ≤-2.5, and neutral if the score above −2.5. [19] And the deleterious prediction was considered for further evaluation. It is available at http://provean.jcvi.org/index.php.

##### 4 Snap2

It is another tool to predict functional effects of mutations. SNAP2 is a trained classifier that is based on a machine learning device called "neural network". It distinguishes between effect and neutral variants/non-synonymous SNPs by taking a variety of sequence and variant features into account. The most important input signal for the prediction is the evolutionary information taken from an automatically generated multiple sequence alignment. Also structural features such as predicted secondary structure and solvent accessibility are considered. If available also annotation (i.e. known functional residues, pattern, regions) of the sequence or close homologs are pulled in. In a cross-validation over 100,000 experimentally annotated variants, SNAP2 reached a sustained two-state accuracy (effect/neutral) of 82% (at an AUC of 0.9). [20] It is available at https://rostlab.org/services/snap2web/.

#### ❖ Tools for Disease related SNPs

##### SNPsGO

Is a web server for predicting disease associated variations from protein sequence and structure. SNPs&GO is an accurate method that, starting from a protein sequence, can predict whether a variation is disease related or not by exploiting the corresponding protein functional annotation. [21] The result obtained from this server consist of three different analytical algorithms; PHD, SNP&GO, and Panther. The output consist of a table listing the number of the mutated position in the protein sequence, the wild-type residue, the new residue and if the related mutation is predicted as disease-related (**Disease** or as neutral polymorphism **Neutral**) and The **RI** value (Reliability Index). It is available at http://snps.biofold.org/snps-and-go/index.html.

##### P-Mut

It is a web-based tool for annotation of pathological variant on proteins. PMut Web portal allows the user to perform pathology predictions, to access a complete repository of pre-calculated predictions, and to generate and validate new predictors. The PMut portal is freely accessible at http://mmb.irbbarcelona.org/PMut. [22]

After using all this analytical algorithms for prediction the result was further analyzed by another tool called I-mutant.

### I-mutant (predictors of effect of single point protein mutation)

It is an online server to predict the protein stability change upon single site mutations starting from protein sequence alone or protein structure when available. It is freely available at (http://gpcr2.biocomp.unibo.it/cgi/predictors/I-Mutant3.0/I-Mutant3.0.cgi). [23]

### Gene mania

it is a free online server that used to predict the function of gene and its interaction using very large set of functional associated data include protein and genetic interactions, pathways, co-expression, co-localization and protein domain similarity.[24] it is available at (http://www.genemania.org/).

### Hope project

It is a webserver that used to analyze the effect of single point mutation on structural level of protein. It is the best way to visualize the mutation since it creates a report consist of figures, animation, 3d structure and mutation report just by submitting the protein sequence and mutation.[25] It is available at http://www.cmbi.ru.nl/hope/.

### Chimera

Ucsf chimera is a computer program that used to visualize the interaction and molecular analysis including density maps, supramolecular assemblies, sequence alignments, docking results, trajectories, and conformational ensembles. It is also allow to create movies. Chimera (version 1.8) software was used to scan the 3D (three-dimensional) structure of specific protein, and then modification was made to the wild type to show the difference after mutation and a graphic view was made for each mutation change [26]. (http://www.cgl.ucsf.edu/chimera/).

## Result

733 SNPs were downloaded from NCBI, from these only 184 SNPs were missense mutations. Firstly we analyzed the effect of the SNPs on the function of the protein using four softwares (SIFT, PROVEAN, Polyhen2, Snap2), resulting in 31 SNPs that had an effect on protein function, then we further analyzed them by SNPs&GO, PMUT and Imutant resulting in 11 SNPs. Finally we studied their effect on the structure of the protein by using Hope and chimera.

**Figure. (1):**
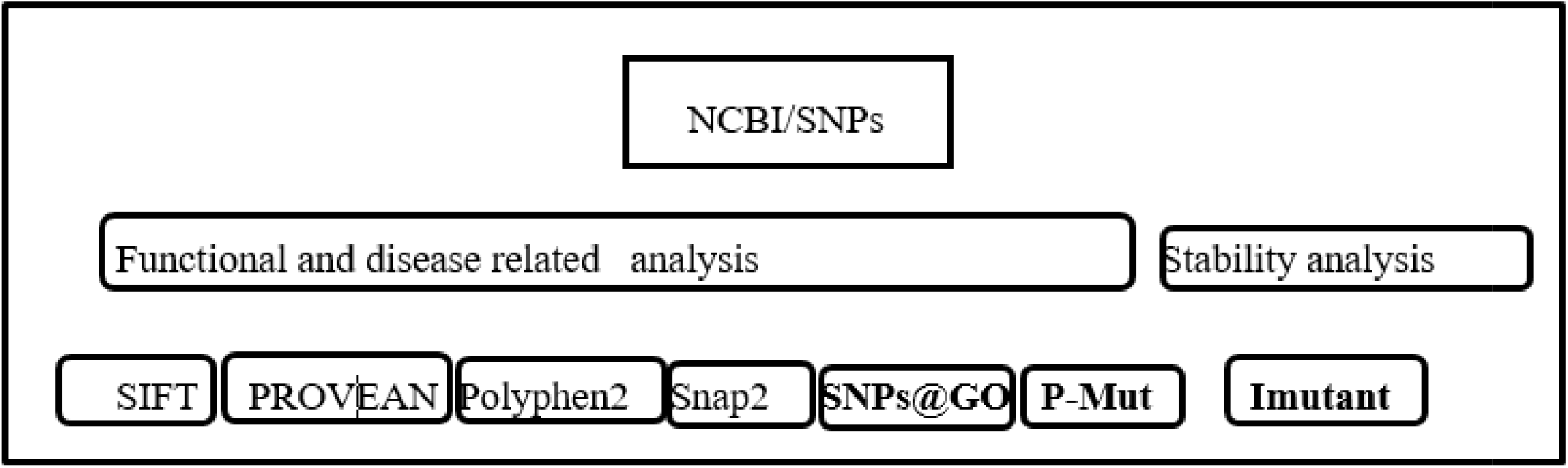
Shows workflow of the paper.

**Table. (1):** SNPs results showing a total of 31 affecting SNPs using SIFT, PROVEAN, PolyPhen-2 and SNAP2 servers.

| dbSNP rs# | sub | sift prediction | sift score | Provean prediction | PROVEAN score | plyphen2 prediction | Polyphen2 score | snap2 prediction | Snap2 Score |
| --- | --- | --- | --- | --- | --- | --- | --- | --- | --- |
| rs28942098 | M1I | affect | 0 | Deleterious | -2.586 | probably damaging | 0.999 | effect | 41 |
| rs1296841626 | L5F | affect | 0 | Deleterious | 2.676 | probably damaging | 1 | effect | 43 |
| rs770544416 | S6G | affect | 0 | Deleterious | -2.539 | possibly damaging | 0.567 | effect | 8 |
| rs1054465259 | G28A | affect | 0 | Deleterious | -3.745 | possibly damaging | 0.816 | effect | 47 |
| rs777541949 | T38S | affect | 0 | Deleterious | -2.785 | probably damaging | 1 | effect | 49 |
| rs1019931450 | Y40C | affect | 0 | Deleterious | -5.114 | probably damaging | 1 | effect | 56 |
| rs200806263 | G49C | affect | 0 | Deleterious | -5.363 | probably damaging | 1 | effect | 14 |
| rs1454615241 | T56I | affect | 0 | Deleterious | -4.102 | probably damaging | 1 | effect | 48 |
| rs1060500015 | L63P | affect | 0 | Deleterious | -4.277 | probably damaging | 1 | effect | 25 |
| rs121434264 | L64P | affect | 0 | Deleterious | -4.435 | probably damaging | 0.999 | effect | 60 |
| rs778467088 | D90H | affect | 0 | Deleterious | -5.36 | probably damaging | 1 | effect | 17 |
| rs1186176634 | S174P | affect | 0 | Deleterious | -2.64 | probably damaging | 1 | effect | 23 |
| rs770439843 | R222G | affect | 0 | Deleterious | -3.272 | possibly damaging | 0.955 | effect | 49 |
| rs776394390 | D223G | affect | 0 | Deleterious | -3.71 | probably damaging | 0.994 | effect | 25 |
| rs1330160847 | I224T | affect | 0 | Deleterious | -3.072 | probably damaging | 0.991 | effect | 25 |
| rs1060500022 | W231R | affect | 0 | Deleterious | -7.888 | probably damaging | 0.996 | effect | 64 |
| rs1452051467 | R234Q | affect | 0 | Deleterious | -3.063 | probably<br>damaging | 1 | effect | 72 |
| rs973863694 | I249T | affect | 0 | Deleterious | -3.648 | probably<br>damaging | 0.999 | effect | 33 |
| rs878855091 | R263C | affect | 0 | Deleterious | -3.402 | probably<br>damaging | 1 | effect | 17 |
| rs1244272523 | R330W | affect | 0 | Deleterious | -3.789 | probably<br>damaging | 0.996 | effect | 56 |
| rs770734388 | P360S | affect | 0 | Deleterious | -6.86 | probably<br>damaging | 1 | effect | 59 |
| rs769288212 | P365L | affect | 0.04 | Deleterious | -8.661 | probably<br>damaging | 1 | effect | 75 |
| rs113200235 | A367V | affect | 0.05 | Deleterious | -3.178 | probably<br>damaging | 0.996 | effect | 66 |
| rs866465727 | V387A | affect | 0 | Deleterious | -3.326 | possibly<br>damaging | 0.656 | effect | 48 |
| rs866793539 | G396C | affect | 0.02 | Deleterious | -7.128 | probably<br>damaging | 1 | effect | 25 |
| rs754454928 | R441C | affect | 0 | Deleterious | -7.186 | probably<br>damaging | 1 | effect | 58 |
| rs778432682 | R441H | affect | 0 | Deleterious | -4.441 | probably<br>damaging | 1 | effect | 64 |
| rs1225502334 | R484C | affect | 0.05 | Deleterious | -3.972 | probably<br>damaging | 0.994 | effect | 32 |
| rs1292596060 | R504S | affect | 0 | Deleterious | -5.296 | probably<br>damaging | 1 | effect | 74 |
| rs759222387 | R504H | affect | 0 | Deleterious | -4.43 | probably<br>damaging | 1 | effect | 73 |
| rs1060500011 | R513W | affect | 0.04 | Deleterious | -3.834 | probably<br>damaging | 0.977 | effect | 59 |

**Table. (2):** SNPs result showing a total of 11 affecting SNPs after using SNPs&GO, PHD and PMUT servers.

| dbSNP<br>rs# | sub | SNPandGO<br>Prediction | RI | snp<br>and go<br>score | PHD<br>Prediction | RI | PHD<br>probability | Pmut<br>prediction | Pmut<br>score |
| --- | --- | --- | --- | --- | --- | --- | --- | --- | --- |
| rs20080<br>6263 | G49<br>C | Disease | 7 | 0.82<br>9 | Disease | 8 | 0.893 | Disease | 0.54<br>(80%) |
| rs10605<br>00015 | L63<br>P | Disease | 3 | 0.65<br>7 | Disease | 7 | 0.861 | Disease | 0.68<br>(85%) |
| rs12143<br>4264 | L64<br>P | Disease | 5 | 0.72<br>9 | Disease | 8 | 0.875 | Disease | 0.75<br>(87%) |
| rs77846<br>7088 | D90<br>H | Disease | 4 | 0.70<br>8 | Disease | 7 | 0.867 | Disease | 0.70<br>(86%) |
| rs77043<br>9843 | R22<br>2G | Disease | 1 | 0.55<br>3 | Disease | 6 | 0.782 | Disease | 0.53<br>(80%) |
| rs10605<br>00022 | W2<br>31R | Disease | 1 | 0.54<br>4 | Disease | 1 | 0.56 | Disease | 0.64<br>(84%) |
| rs77073<br>4388 | P36<br>0S | Disease | 3 | 0.65 | Disease | 0 | 0.503 | Disease | 0.84<br>(90%) |
| rs75445<br>4928 | R44<br>1C | Disease | 5 | 0.74<br>3 | Disease | 5 | 0.775 | Disease | 0.61<br>(83%) |
| rs77843<br>2682 | R44<br>1H | Disease | 3 | 0.66<br>9 | Disease | 5 | 0.753 | Disease | 0.70<br>(86%) |
| rs12925<br>96060 | R50<br>4S | Disease | 6 | 0.82<br>3 | Disease | 7 | 0.867 | Disease | 0.86<br>(91%) |
| rs75922<br>2387 | R50<br>4H | Disease | 6 | 0.81<br>1 | Disease | 7 | 0.857 | Disease | 0.72<br>(86%) |

**Table. (3):**
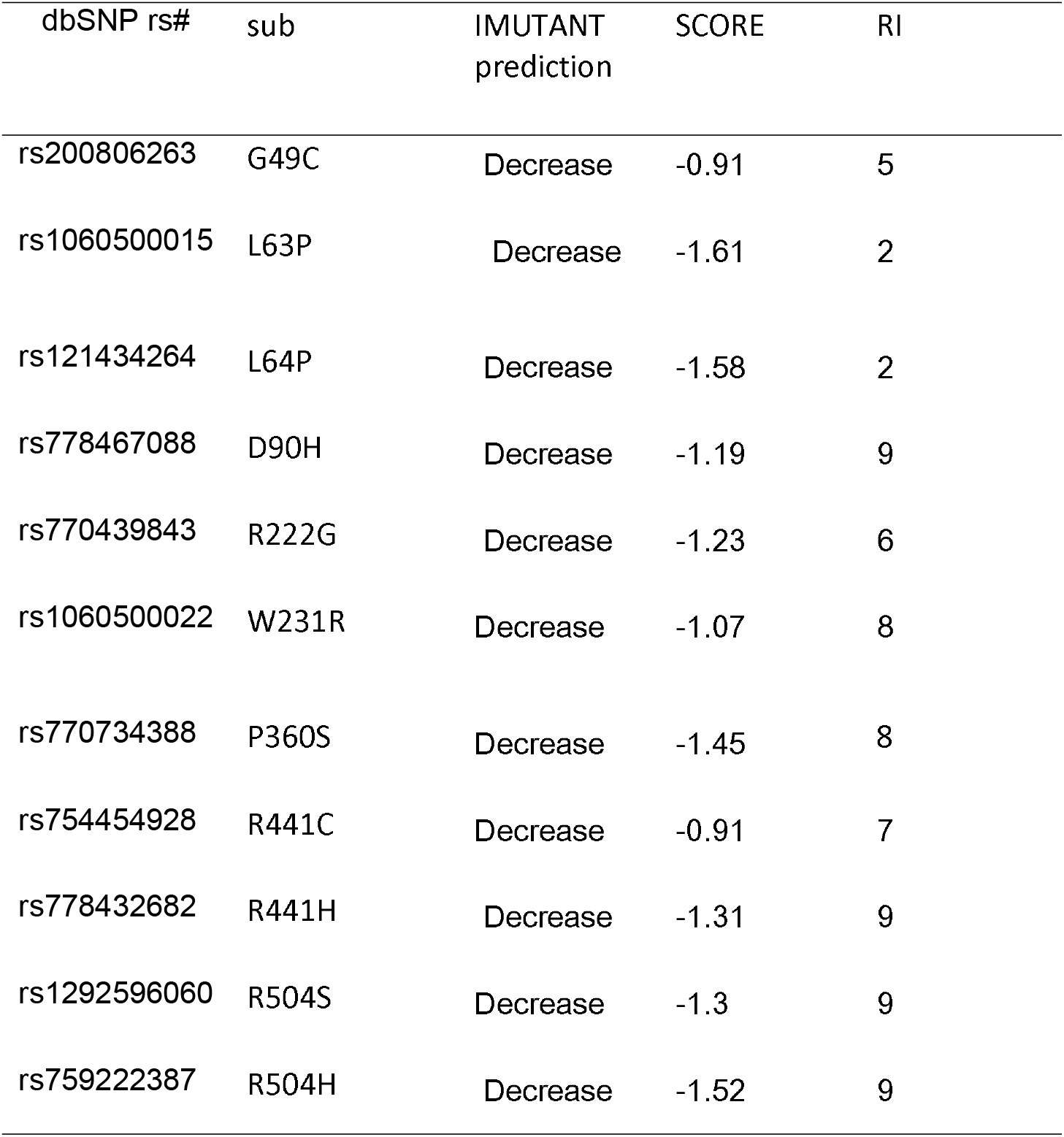
Stability analysis for the 11 affecting SNPs using I-Mutant server.

**Table. (4):** *CDC73* gene Functions and its appearance in network and genome.

| Function | FDR | Genes in network | Genes in genome |
| --- | --- | --- | --- |
| transcription elongation factor complex | 2.52E-09 | 6 | 28 |
| regulation of transcription elongation from RNA polymerase II promoter | 2.52E-09 | 5 | 10 |
| transcription elongation from RNA polymerase II promoter | 6.60E-09 | 7 | 75 |
| DNA-templated transcription, elongation | 5.45E-08 | 7 | 108 |
| positive regulation of DNA-templated transcription, elongation | 5.45E-08 | 5 | 20 |
| mRNA polyadenylation | 7.21E-08 | 5 | 22 |
| RNA polyadenylation | 9.97E-08 | 5 | 24 |
| regulation of DNA-templated transcription, elongation | 2.01E-07 | 5 | 28 |
| histone modification | 3.31E-07 | 8 | 260 |
| covalent chromatin modification | 3.47E-07 | 8 | 265 |
| DNA-directed RNA polymerase II, holoenzyme | 3.50E-07 | 6 | 81 |
| endodermal cell fate commitment | 3.50E-07 | 4 | 10 |
| RNA polymerase complex | 7.92E-07 | 6 | 96 |
| DNA-directed RNA polymerase complex | 7.92E-07 | 6 | 95 |
| nuclear DNA-directed RNA polymerase complex | 7.92E-07 | 6 | 95 |
| endodermal cell differentiation | 8.50E-07 | 4 | 13 |
| negative regulation of myeloid cell differentiation | 1.03424E-06 | 5 | 44 |
| cell fate commitment involved in formation of primary germ layer | 2.50852E-06 | 4 | 17 |
| histone monoubiquitination | 3.86491E-06 | 4 | 19 |
| endoderm formation | 5.6616E-06 | 4 | 21 |
| mRNA processing | 1.22928E-05 | 7 | 287 |
| endoderm development | 1.28119E-05 | 4 | 26 |
| histone ubiquitination | 2.24022E-05 | 4 | 30 |
| mRNA 3'-end processing | 2.5645E-05 | 5 | 88 |
| regulation of myeloid cell differentiation | 4.02517E-05 | 5 | 97 |
| protein monoubiquitination | 4.41323E-05 | 4 | 37 |
| formation of primary germ layer | 4.41323E-05 | 4 | 37 |
| RNA 3'-end processing | 4.41323E-05 | 5 | 101 |
| regulation of mRNA processing | 0.000289177 | 4 | 59 |
| positive regulation of mRNA 3'-end processing | 0.000313172 | 3 | 15 |
| regulation of mRNA 3'-end processing | 0.000372752 | 3 | 16 |
| myeloid cell differentiation | 0.000446549 | 5 | 165 |
| Gastrulation | 0.000451884 | 4 | 68 |
| positive regulation of mRNA processing | 0.000804394 | 3 | 21 |
| histone H3-K4 methylation | 0.001915467 | 3 | 28 |
| cell fate commitment | 0.002956445 | 4 | 111 |
| negative regulation of cell differentiation | 0.004090933 | 5 | 267 |
| stem cell maintenance | 0.005276884 | 3 | 40 |
| histone lysine methylation | 0.006873226 | 3 | 44 |
| regulation of histone modification | 0.011780086 | 3 | 53 |
| histone methylation | 0.01357129 | 3 | 56 |
| stem cell differentiation | 0.013955822 | 4 | 171 |
| cellular response to lipopolysaccharide | 0.01758198 | 3 | 62 |
| regulation of chromatin organization | 0.018029071 | 3 | 63 |
| cellular response to molecule of bacterial origin | 0.020035617 | 3 | 66 |
| embryonic morphogenesis | 0.020035617 | 4 | 192 |
| RNA polymerase II core binding | 0.020433712 | 2 | 10 |
| peptidyl-lysine trimethylation | 0.022132906 | 2 | 11 |
| basal transcription machinery binding | 0.022132906 | 2 | 11 |
| basal RNA polymerase II transcription machinery binding | 0.022132906 | 2 | 11 |
| mRNA cleavage | 0.022132906 | 2 | 11 |
| transcriptionally active chromatin | 0.022132906 | 2 | 11 |
| regulation of histone H3-K4 methylation | 0.022132906 | 2 | 11 |
| protein alkylation | 0.022434688 | 3 | 73 |
| protein methylation | 0.022434688 | 3 | 73 |
| cellular response to biotic stimulus | 0.022949254 | 3 | 74 |
| mRNA cleavage factor complex | 0.024254179 | 2 | 12 |
| RNA polymerase core enzyme binding | 0.024254179 | 2 | 12 |
| positive regulation of histone methylation | 0.028159975 | 2 | 13 |
| response to lipopolysaccharide | 0.037153654 | 3 | 90 |
| stem cell development | 0.037153654 | 3 | 90 |
| regulation of chromosome organization | 0.037153654 | 3 | 90 |
| RNA polymerase binding | 0.040493707 | 2 | 16 |
| regulation of histone methylation | 0.04514658 | 2 | 17 |
| response to molecule of bacterial origin | 0.049891593 | 3 | 101 |
| peptidyl-lysine methylation | 0.054973868 | 2 | 19 |
| macromolecule methylation | 0.080506456 | 3 | 120 |

**\*FDR:** false discovery rate is greater than or equal to the probability that this is a false positive.

**Table. (5):** The gene co-expressed, share domain and interaction with *CDC73* gene network.

**\*FDR:** false discovery rate is greater than or equal to the probability that this is a false positive. **Table. (5):**The gene co-expressed, share domain and interaction with *CDC73* gene network.
| Gene 1 | Gene 2 | Weight | Network group |
| --- | --- | --- | --- |
| AP3S1 | HSP90AA1 | 0.013259681 | Co-expression |
| CTR9 | CDC73 | 0.008208696 | Co-expression |
| CTR9 | LEO1 | 0.003100712 | Co-expression |
| WDR61 | LEO1 | 0.007506342 | Co-expression |
| WDR61 | CTR9 | 0.003934787 | Co-expression |
| KMT2C | BCL9L | 0.01605392 | Co-expression |
| CDK9 | CPSF4 | 0.007431932 | Co-expression |
| GTF2F1 | PAF1 | 0.004063675 | Co-expression |
| GTF2F1 | KMT2C | 0.008976047 | Co-expression |
| AURKB | TKT | 0.011608324 | Co-expression |
| NUP98 | CTR9 | 0.014227687 | Co-expression |
| CPSF4 | TKT | 0.00430011 | Co-expression |
| CPSF4 | AURKB | 0.006637127 | Co-expression |
| CHUK | CSTF3 | 0.016616315 | Co-expression |
| CSTF2 | AURKB | 0.004579851 | Co-expression |
| CHUK | CDC73 | 0.001583963 | Co-expression |
| CHUK | NUP98 | 0.004846978 | Co-expression |
| GTF2F1 | TKT | 0.009125041 | Co-expression |
| CPSF4 | CSTF3 | 0.014529244 | Co-expression |
| CHUK | NUP98 | 0.008912819 | Co-expression |
| NUP98 | CDC73 | 0.00588025 | Co-expression |
| CHUK | CDC73 | 0.001885684 | Co-expression |
| CSTF2 | LEO1 | 0.006971632 | Co-expression |
| CSTF3 | CTR9 | 0.006781118 | Co-expression |
| CHUK | CDC73 | 0.011535326 | Co-expression |
| CHUK | CTR9 | 0.00340099 | Co-expression |
| AP3S1 | CTR9 | 0.004597292 | Co-expression |
| GTF2F1 | PAF1 | 0.010657921 | Co-expression |
| GTF2F1 | CPSF4 | 0.011196721 | Co-expression |
| CSTF2 | CTR9 | 0.011377413 | Co-expression |
| CDK9 | CPSF4 | 0.009146069 | Co-expression |
| AURKB | CSTF3 | 0.006553773 | Co-expression |
| SPCS3 | CHUK | 0.00718225 | Co-expression |
| AURKB | NUP98 | 0.004425747 | Pathway |
| CPSF4 | CSTF3 | 0.11179613 | Pathway |
| CHUK | HSP90AA1 | 0.006348364 | Pathway |
| CSTF2 | CSTF3 | 0.009734826 | Pathway |
| CSTF2 | CPSF4 | 0.11179613 | Pathway |
| GTF2F1 | CSTF3 | 0.005905584 | Pathway |
| GTF2F1 | CSTF2 | 0.005905584 | Pathway |
| GTF2F1 | CDK9 | 0.011108032 | Pathway |
| CSTF2 | CSTF3 | 0.011789562 | Pathway |
| GTF2F1 | CSTF3 | 0.008021638 | Pathway |
| GTF2F1 | CSTF2 | 0.007878398 | Pathway |
| GTF2F1 | CDK9 | 0.014240757 | Pathway |
| CHUK | HSP90AA1 | 0.088913664 | Pathway |
| LEO1 | CDC73 | 0.122442566 | Physical Interactions |
| PAF1 | CDC73 | 0.21769759 | Physical Interactions |
| PAF1 | LEO1 | 0.15407903 | Physical Interactions |
| CTR9 | CDC73 | 0.21769759 | Physical Interactions |
| CTR9 | LEO1 | 0.15407903 | Physical Interactions |
| CTR9 | PAF1 | 0.27394587 | Physical Interactions |
| WDR61 | CDC73 | 0.21769759 | Physical Interactions |
| WDR61 | LEO1 | 0.15407903 | Physical Interactions |
| WDR61 | PAF1 | 0.27394587 | Physical Interactions |
| WDR61 | CTR9 | 0.27394587 | Physical Interactions |
| HSP90AA1 | CDC73 | 0.3958806 | Physical Interactions |
| CDK9 | HSP90AA1 | 0.20299108 | Physical Interactions |
| CDK9 | CDC73 | 0.119673245 | Physical Interactions |
| AURKB | CDC73 | 0.1827706 | Physical Interactions |
| AURKB | HSP90AA1 | 0.042777777 | Physical Interactions |
| PAF1 | CDC73 | 0.13507365 | Physical Interactions |
| CTR9 | CDC73 | 0.18643428 | Physical Interactions |
| NUP98 | CDC73 | 0.203537 | Physical Interactions |
| NUP98 | PAF1 | 0.31207067 | Physical Interactions |
| CHUK | CDC73 | 0.1774874 | Physical Interactions |
| CHUK | PAF1 | 0.2721304 | Physical Interactions |
| CDK9 | HSP90AA1 | 0.040214784 | Physical Interactions |
| LEO1 | CDC73 | 0.20954604 | Physical Interactions |
| PAF1 | CDC73 | 0.16041815 | Physical Interactions |
| PAF1 | LEO1 | 0.08672058 | Physical Interactions |
| CTR9 | CDC73 | 0.24223034 | Physical Interactions |
| CTR9 | LEO1 | 0.1309475 | Physical Interactions |
| CTR9 | PAF1 | 0.10024697 | Physical Interactions |
| CSTF2 | CSTF3 | 0.24596581 | Physical Interactions |
| GTF2F1 | CDC73 | 0.1822143 | Physical Interactions |
| GTF2F1 | CDK9 | 0.077137925 | Physical Interactions |
| LEO1 | CDC73 | 0.3225924 | Physical Interactions |
| PAF1 | CDC73 | 0.3225924 | Physical Interactions |
| CTR9 | CDC73 | 0.3225924 | Physical Interactions |
| WDR61 | CDC73 | 0.17647609 | Physical Interactions |
| CSTF3 | CDC73 | 0.3225924 | Physical Interactions |
| FIP1L1 | CDC73 | 0.3225924 | Physical Interactions |
| CPSF4 | CDC73 | 0.17647609 | Physical Interactions |
| CHUK | HSP90AA1 | 0.02887361 | Physical Interactions |
| KMT2C | CDC73 | 0.15483053 | Physical Interactions |
| CSTF2 | CDC73 | 0.16203262 | Physical Interactions |
| LEO1 | CDC73 | 0.09317254 | Physical Interactions |
| PAF1 | CDC73 | 0.08377836 | Physical Interactions |
| PAF1 | LEO1 | 0.07275753 | Physical Interactions |
| CTR9 | CDC73 | 0.09010142 | Physical Interactions |
| CTR9 | LEO1 | 0.078248814 | Physical Interactions |
| CTR9 | PAF1 | 0.07035932 | Physical Interactions |
| WDR61 | CDC73 | 0.114276804 | Physical Interactions |
| WDR61 | LEO1 | 0.09924398 | Physical Interactions |
| WDR61 | PAF1 | 0.089237645 | Physical Interactions |
| WDR61 | CTR9 | 0.09597274 | Physical Interactions |
| AP3S1 | CDC73 | 0.2352175 | Physical Interactions |
| CSTF2 | CSTF3 | 0.077111326 | Physical Interactions |
| LEO1 | CDC73 | 0.049589828 | Physical Interactions |
| PAF1 | CDC73 | 0.10079521 | Physical Interactions |
| PAF1 | LEO1 | 0.11382505 | Physical Interactions |
| CTR9 | CDC73 | 0.01533186 | Physical Interactions |
| CTR9 | LEO1 | 0.017313818 | Physical Interactions |
| CTR9 | PAF1 | 0.035191692 | Physical Interactions |
| WDR61 | CDC73 | 0.041940387 | Physical Interactions |
| WDR61 | LEO1 | 0.047362044 | Physical Interactions |
| WDR61 | PAF1 | 0.09626707 | Physical Interactions |
| WDR61 | CTR9 | 0.014643089 | Physical Interactions |
| FIP1L1 | CSTF3 | 0.113308094 | Physical Interactions |
| CPSF4 | FIP1L1 | 0.14965165 | Physical Interactions |
| CSTF2 | FIP1L1 | 0.10615889 | Physical Interactions |
| CDK9 | CTR9 | 0.0411037 | Physical Interactions |
| LEO1 | CDC73 | 0.05172406 | Physical Interactions |
| PAF1 | CDC73 | 0.028004196 | Physical Interactions |
| PAF1 | LEO1 | 0.054951686 | Physical Interactions |
| CTR9 | CDC73 | 0.04297603 | Physical Interactions |
| CTR9 | LEO1 | 0.08433042 | Physical Interactions |
| CTR9 | PAF1 | 0.045657773 | Physical Interactions |
| WDR61 | CDC73 | 0.04966663 | Physical Interactions |
| WDR61 | LEO1 | 0.09745915 | Physical Interactions |
| WDR61 | PAF1 | 0.05276587 | Physical Interactions |
| WDR61 | CTR9 | 0.080975994 | Physical Interactions |
| CSTF3 | CDC73 | 0.06669109 | Physical Interactions |
| FIP1L1 | CSTF3 | 0.14174445 | Physical Interactions |
| AURKB | HSP90AA1 | 0.002030884 | Physical Interactions |
| CPSF4 | CDC73 | 0.09574746 | Physical Interactions |
| CPSF4 | FIP1L1 | 0.20350051 | Physical Interactions |
| CHUK | HSP90AA1 | 0.001720827 | Physical Interactions |
| BCL9L | CDC73 | 0.2791709 | Physical Interactions |
| KMT2C | CDC73 | 0.04642578 | Physical Interactions |
| CSTF2 | CDC73 | 0.0463337 | Physical Interactions |
| CSTF2 | CSTF3 | 0.11722768 | Physical Interactions |
| CDK9 | PAF1 | 0.01438359 | Physical Interactions |
| GTF2F1 | CDK9 | 0.013446528 | Physical Interactions |
| AURKB | HSP90AA1 | 0.03819712 | Physical Interactions |
| CDK9 | HSP90AA1 | 0.03819712 | Physical Interactions |
| LEO1 | CDC73 | 0.5193767 | Physical Interactions |
| PAF1 | CDC73 | 0.5193767 | Physical Interactions |
| CTR9 | CDC73 | 0.37205398 | Physical Interactions |
| WDR61 | CDC73 | 0.5193767 | Physical Interactions |
| AURKB | HSP90AA1 | 0.080966905 | Physical Interactions |
| CDK9 | HSP90AA1 | 0.036444858 | Physical Interactions |
| LEO1 | CDC73 | 0.0771831 | Physical Interactions |
| PAF1 | CDC73 | 0.031383026 | Physical Interactions |
| PAF1 | LEO1 | 0.14005615 | Physical Interactions |
| CTR9 | CDC73 | 0.058806863 | Physical Interactions |
| CTR9 | PAF1 | 0.10671071 | Physical Interactions |
| WDR61 | CTR9 | 0.34459004 | Physical Interactions |
| CPSF4 | FIP1L1 | 0.44817418 | Physical Interactions |
| CHUK | HSP90AA1 | 0.0081622 | Physical Interactions |
| HSF2BP | CDC73 | 0.31519976 | Physical Interactions |
| CDK9 | HSP90AA1 | 0.010792454 | Physical Interactions |
| HSF2BP | CDC73 | 0.58701295 | Physical Interactions |
| CPSF4 | FIP1L1 | 0.6677753 | Physical Interactions |
| CHUK | HSP90AA1 | 0.007963367 | Physical Interactions |
| CSTF2 | CSTF3 | 0.2417784 | Physical Interactions |
| CTR9 | LEO1 | 0.09094542 | Predicted |
| CTR9 | LEO1 | 0.0869592 | Predicted |
| LEO1 | CDC73 | 1 | Predicted |
| CPSF4 | FIP1L1 | 1 | Predicted |
| CSTF3 | LEO1 | 0.020077549 | Predicted |
| FIP1L1 | CSTF3 | 0.015798066 | Predicted |
| NUP98 | CTR9 | 0.047920566 | Predicted |
| SPCS3 | CDC73 | 0.25866398 | Predicted |
| CTR9 | LEO1 | 0.34126943 | Predicted |
| TKT | CDC73 | 1 | Predicted |
| TKT | CDC73 | 1 | Predicted |
| CHUK | AURKB | 0.006874138 | Shared protein domains |
| CDK9 | AURKB | 0.003942981 | Shared protein domains |
| CDK9 | CHUK | 0.006889931 | Shared protein domains |
| CHUK | AURKB | 0.003882364 | Shared protein domains |
| CDK9 | AURKB | 0.002896895 | Shared protein domains |
| CDK9 | CHUK | 0.004389529 | Shared protein domains |

**Figure. (2):**
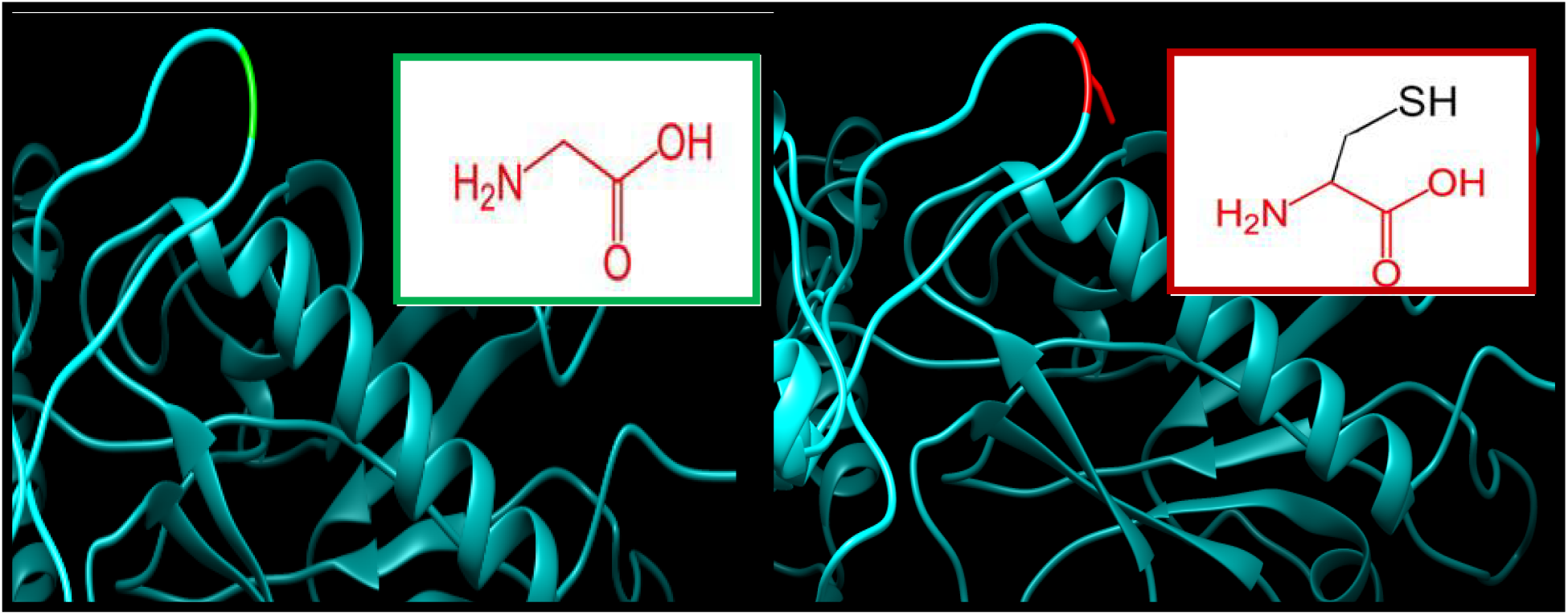
G49C: Glycine changed to Cysteine at position 49.

**Figure. (3):**
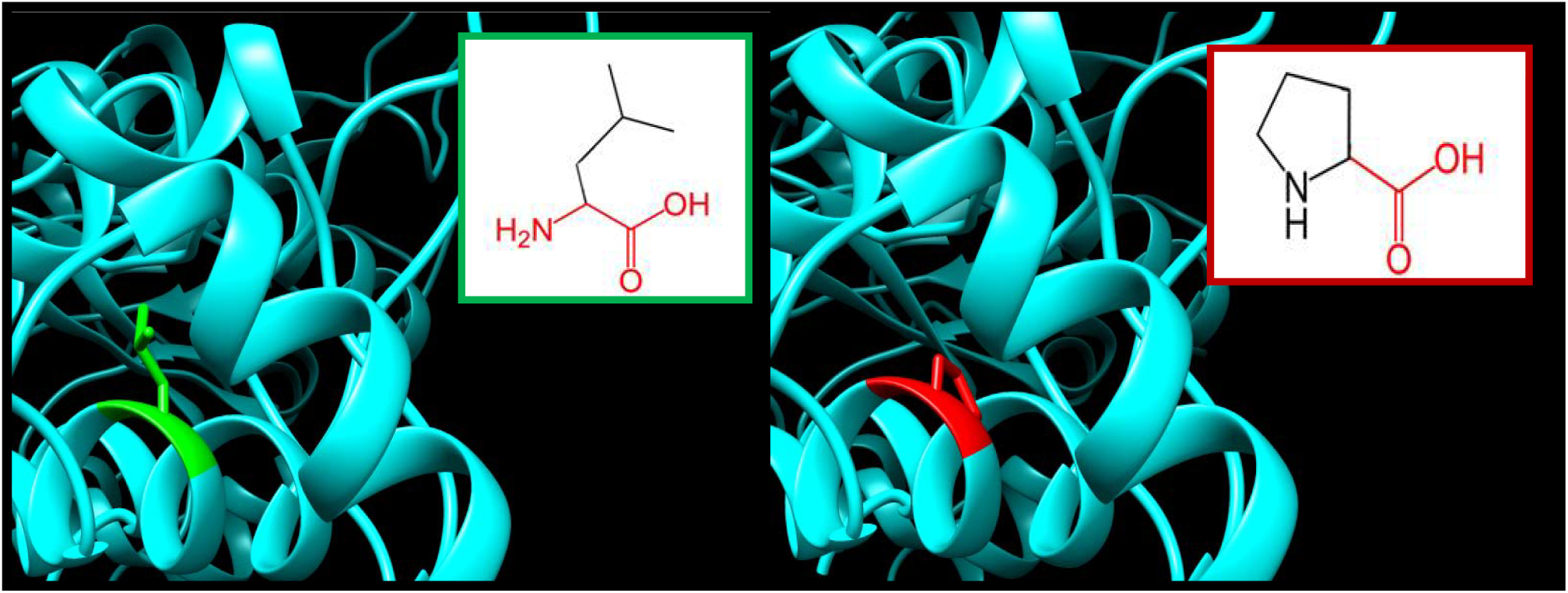
L63P: Leucine changed to Proline at position 63.

**Figure. (4):**
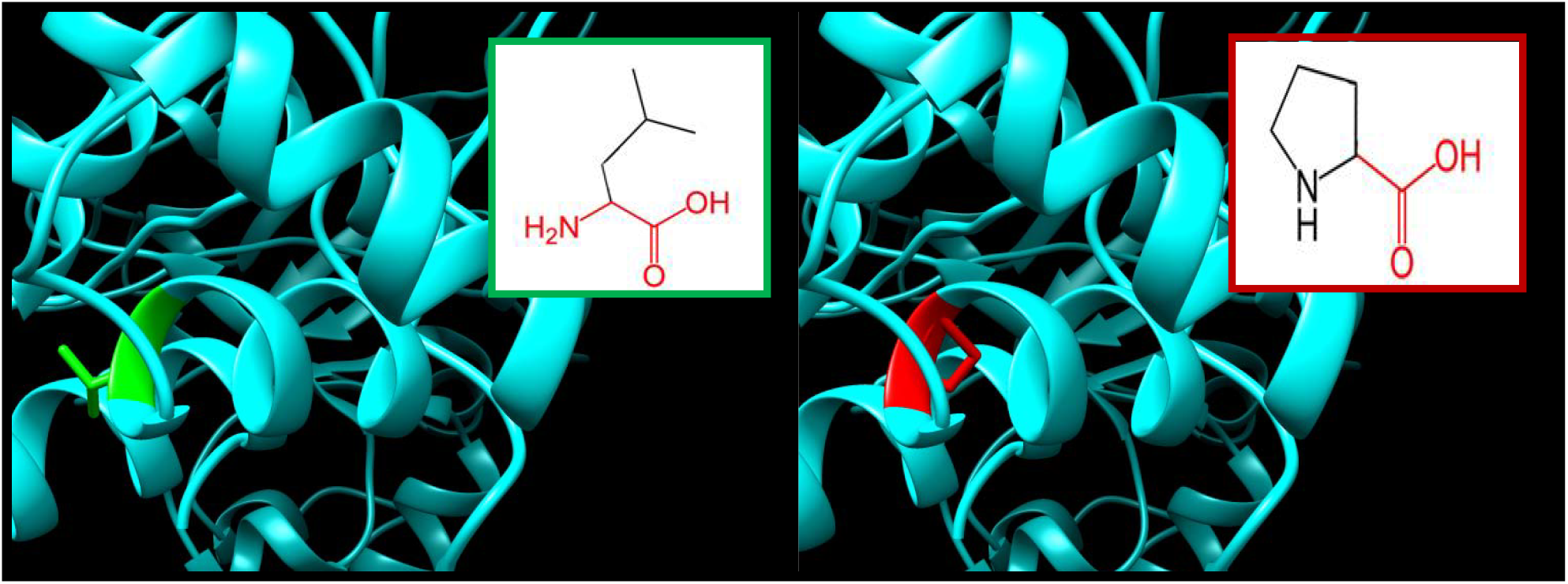
L64P: Leucine to Proline at position 64.

**Figure. (5):**
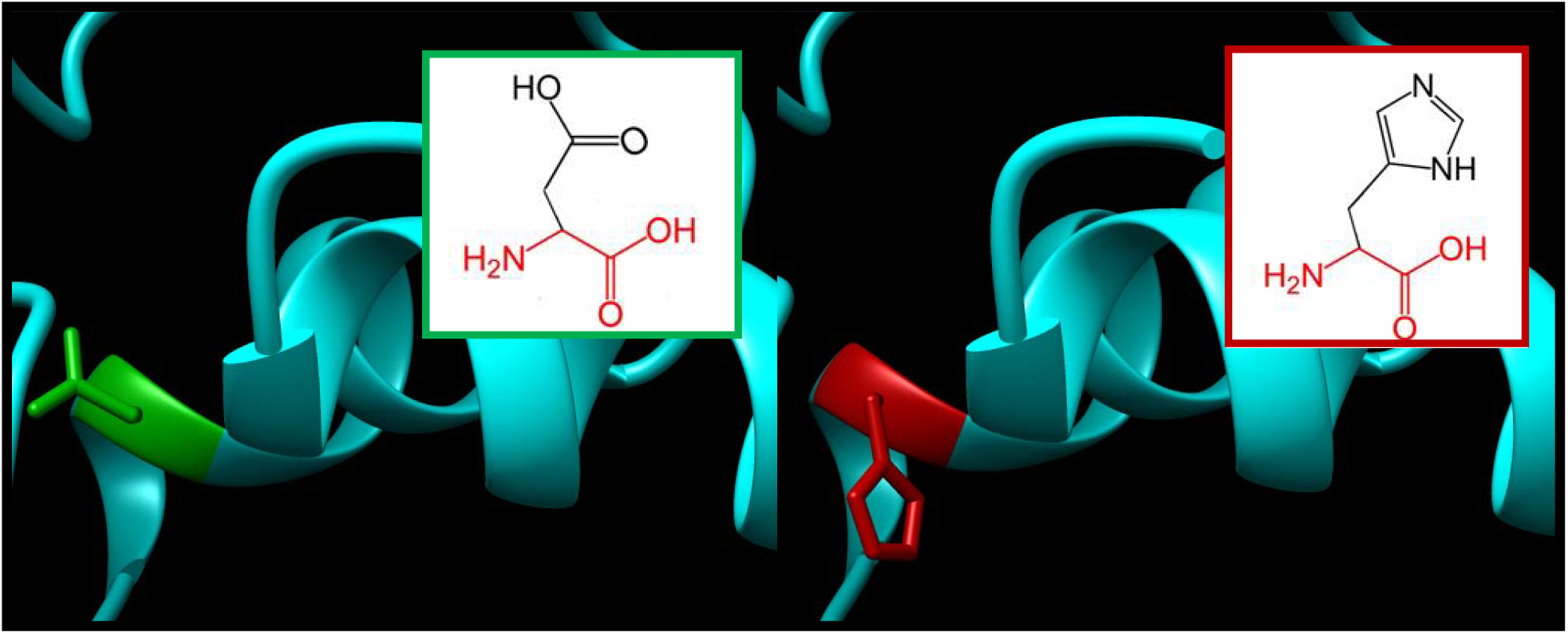
D90H: Aspartic acid changed to Histidine at position 90.

**Figure. (6):**
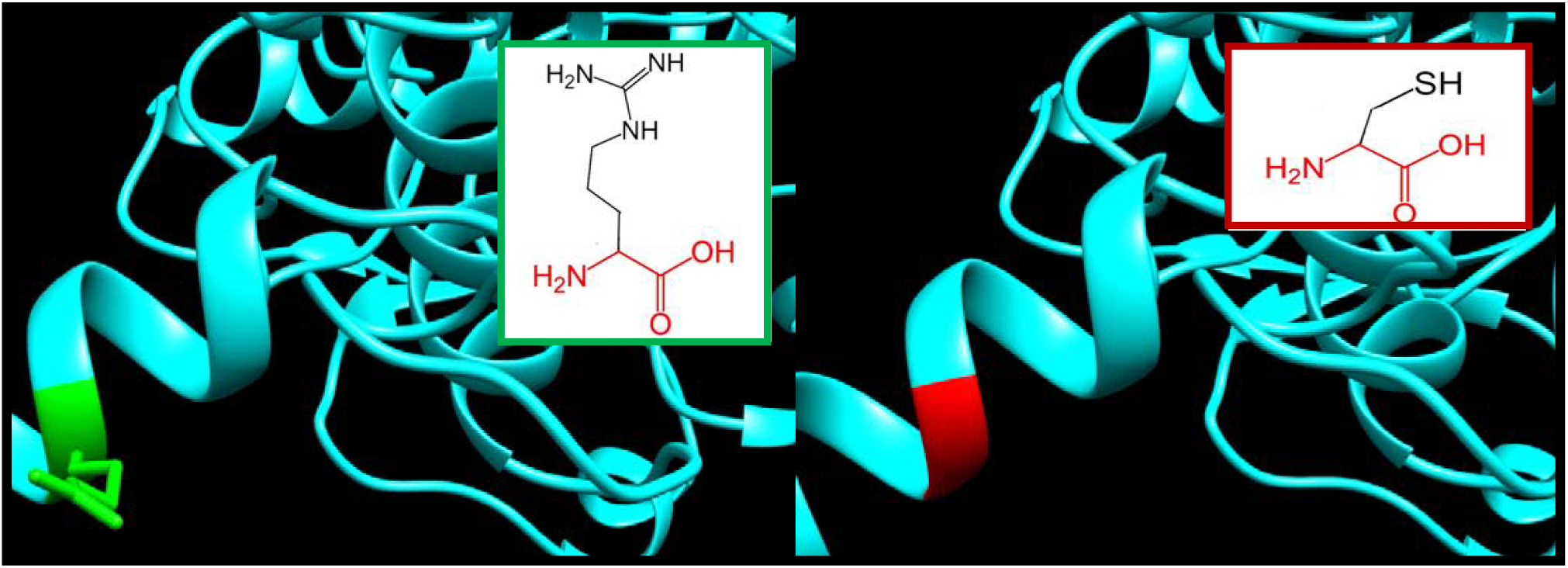
R222G: Arginine changed to Glycine at position 222.

**Figure. (7):**
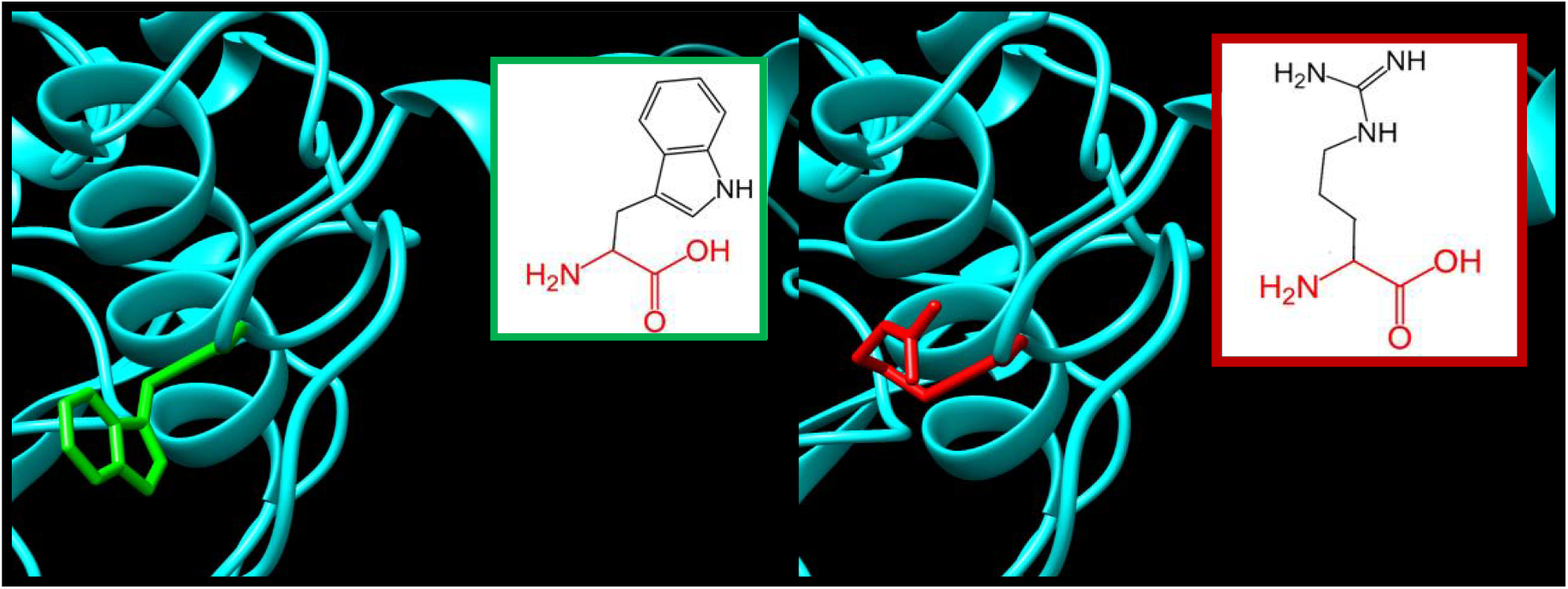
W231R: Tryptophan changed to Arginine at position 231.

**Figure. (8):**
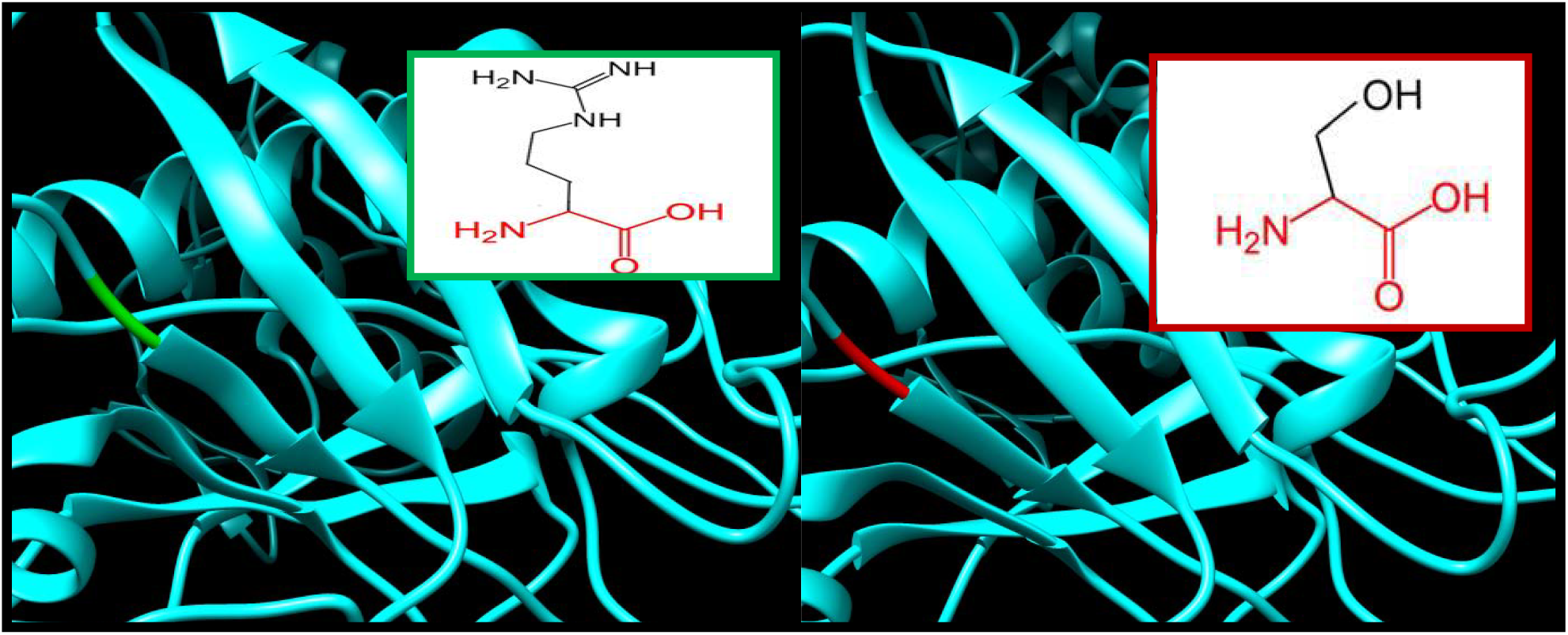
P360S: Proline changed to Serine at position 360.

**Figure. (9):**
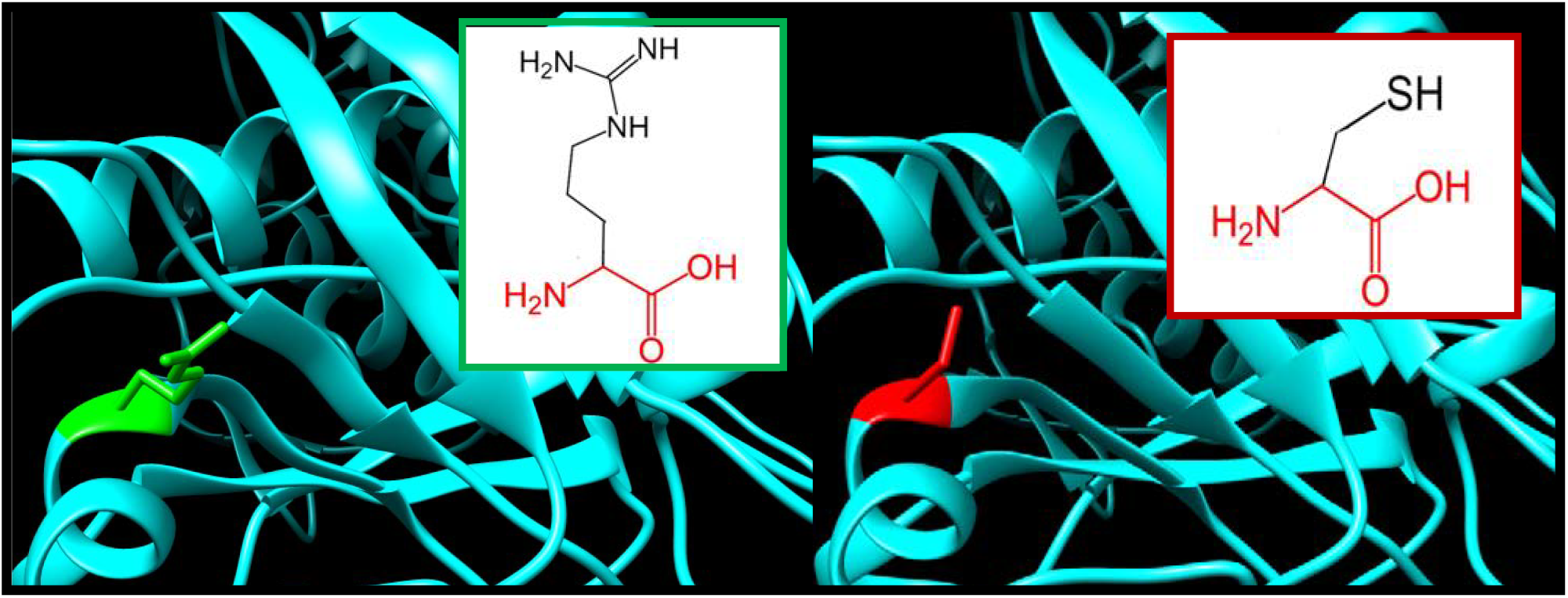
R441C: Arginine changed to Cysteine at position 441.

**Figure. (10):**
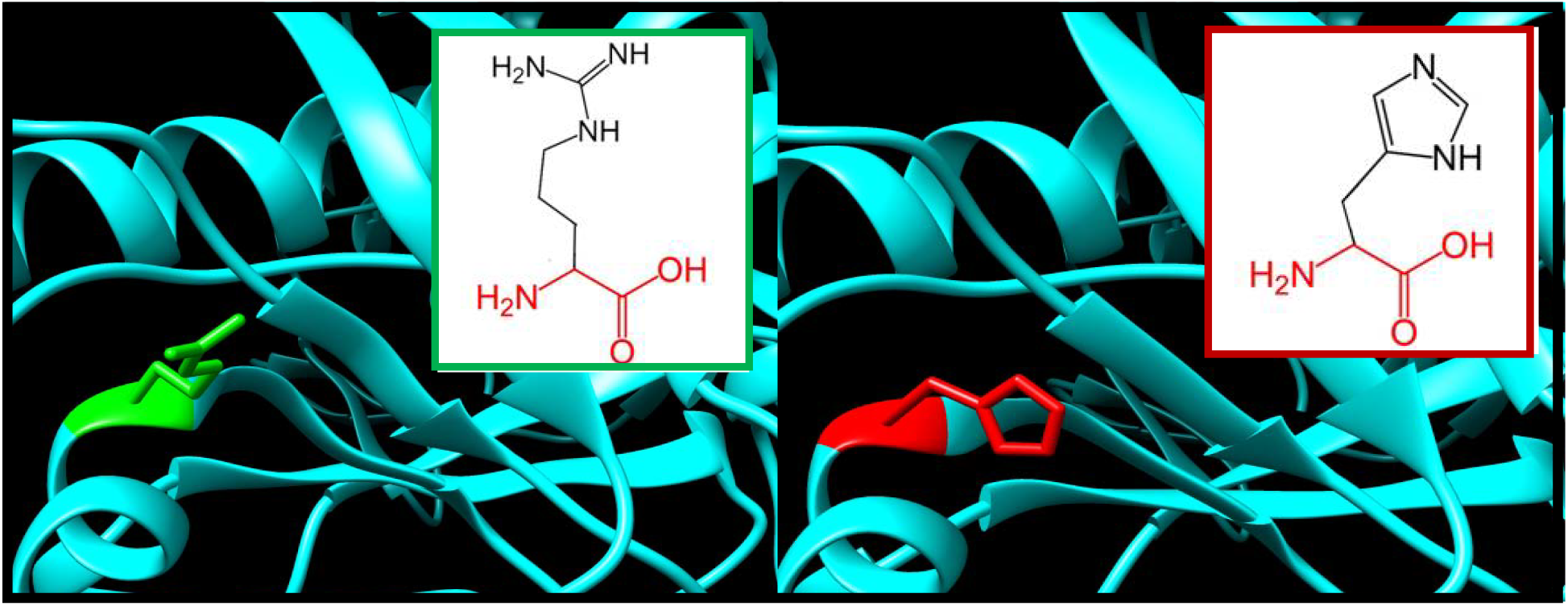
R441H: Arginine changed to Histidine at position 441.

**Figure. (11):**
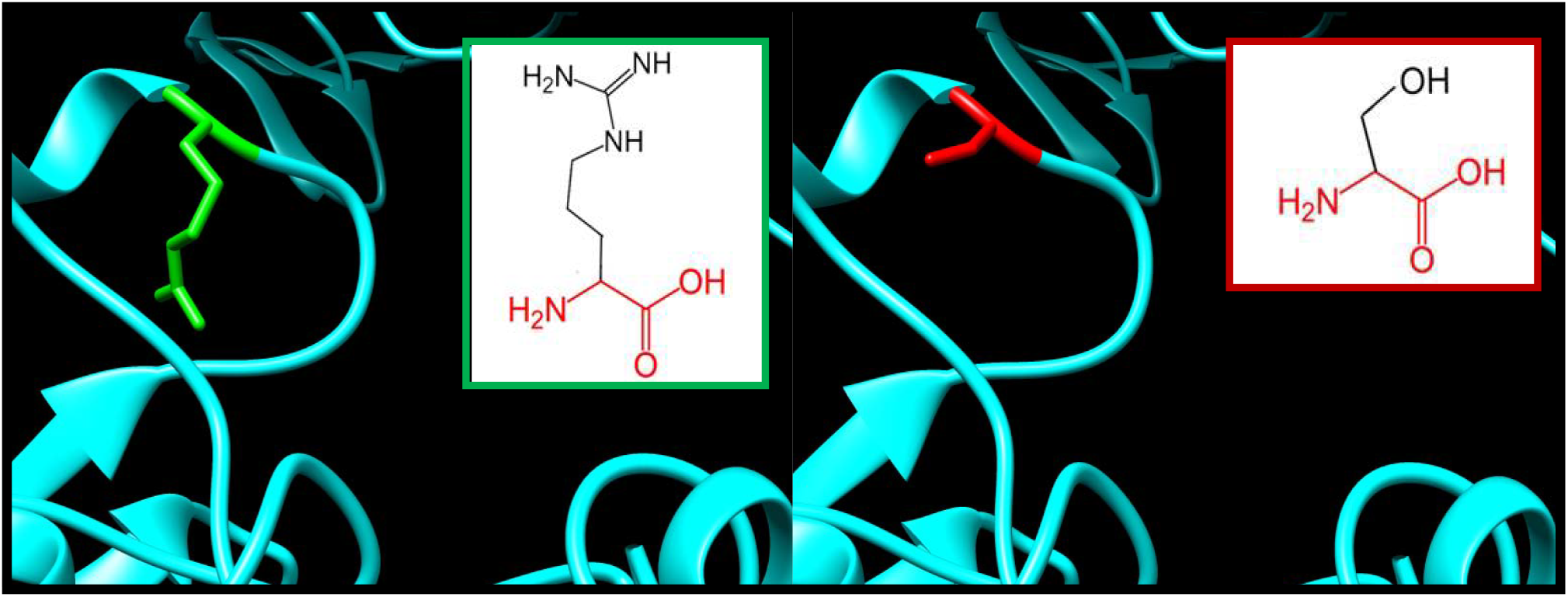
R504S: Arginine changed to Serine at position 504.

**Figure. (12):**
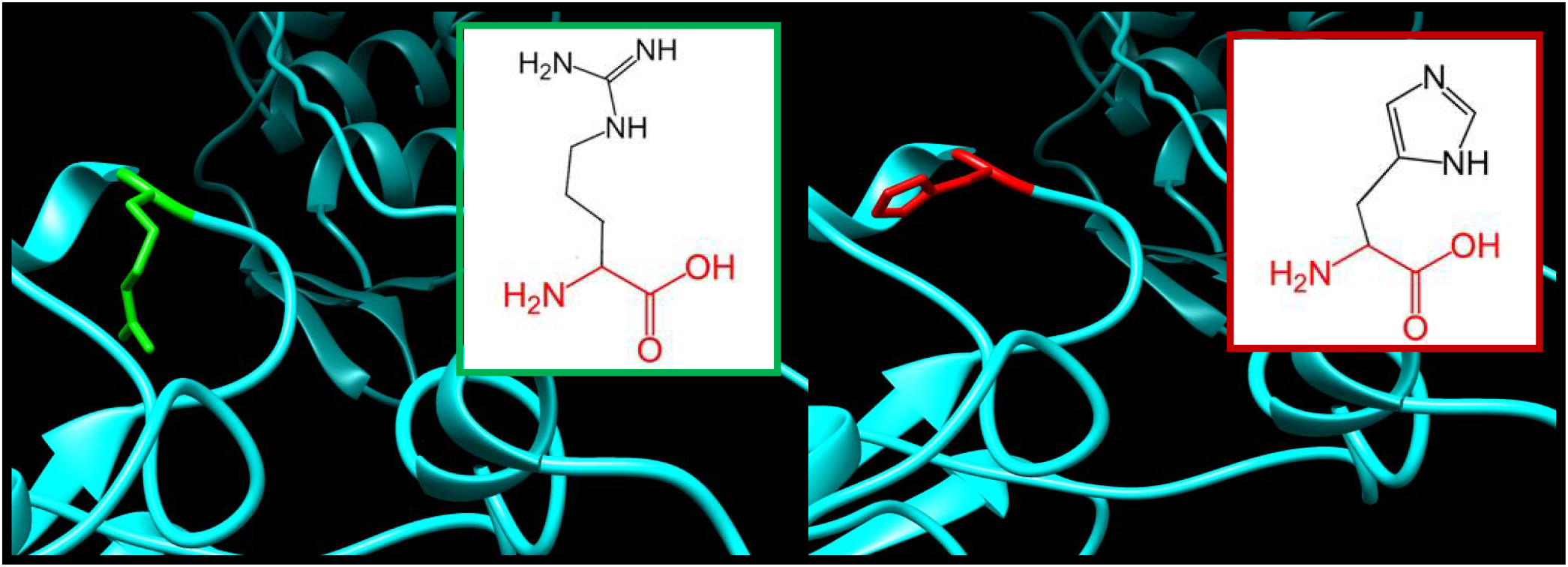
R504H: Arginine changed to Histidine at position 504.

**Figure. (13):**
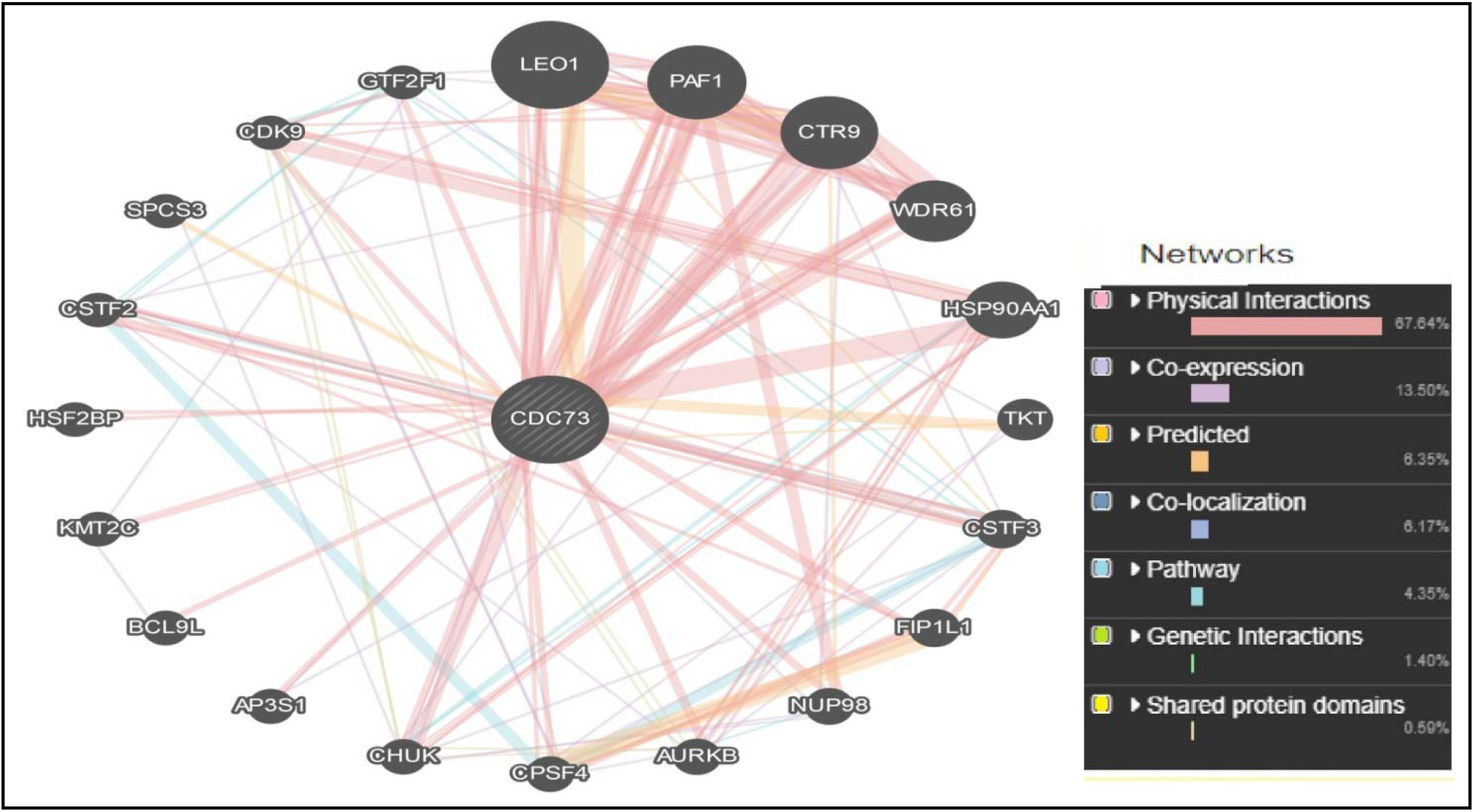
Shows interaction between *CDC73* gene and related genes using GeneMANIA.

## Discussion

11 novel mutations were found to have damaging impact on the structure and function of the protein, out of 184 missense SNPs download from NCBI web side. The SNPs downloaded were analyzed using eight softwares to study the effect of the mutation on the structural and function of the protein.

Project hope was used to the study the effect of the mutation on the physiochemical properties of the protein, where we have found changes on the charge, size and hydrophobicity of the protein illustrated in the hope report. We also recognized a common domains which were Cdc73/Parafibromin IPR007852, Cell Division Control Protein 73, C-Terminal Domain Superfamily IPR038103 and Cell Division Control Protein 73, C-Terminal IPR031336, which indicated the conservancy and the significance of these SNPs.

Chimera software was used to show the 3D changes in the structure of the protein coupled with schematic structures from project hope for comparison, which further prove the effect of these SNPs on the structure of the protein.

Previous papers report novel mutations of deletions type in c.191-192 delT, [27] and (c.1379delT/p.L460Lfs*18). [28] While novel deletion of exons 4 to 10 of CDC73 was detected in another study. [29]

In other study NGS revealed four pathogenic or likely pathogenic germline sequence variants in *CDC73* c.271C>T (p.Arg91*), c.496C>T (p.Gln166*), c.685A>T (p.Arg229*) and c.787C>T (p.Arg263Cys). [2]

This study identified 11 novel mutations in *CDC73* gene related to jaw syndrome which could be used as diagnostic marker for the disease, and also serve as “actionable targets” for chemotherapeutic intervention in patients whose disease is no longer surgically curable Further in vitro and in vivo studies are needed to confirm these results.

## Conclusion

In this study the effect of the SNPs of *CDC73* gene was thoroughly investigated through different bioinformatics prediction softwares.11 novel mutations were found to have damaging impact on the structure and function of the protein and may thus be used as diagnostic marker for the disease.

## Acknowledgement

The authors wish to acknowledge the enthusiastic cooperation of Africa City of Technology - Sudan.

